# Analysis of the blood microbiome in a porcine model of fecal-induced peritonitis

**DOI:** 10.1101/2021.02.18.431914

**Authors:** Hwi Hyun, Min Seok Lee, Inwon Park, Hwa Soo Ko, Seungmin Yun, Dong-Hyun Jang, Seonghye Kim, Hajin Kim, Joo H. Kang, Jae Hyuk Lee, Taejoon Kwon

## Abstract

**Background:** Recent studies have proposed the existence of a blood microbiome, even in the healthy host. However, we do not know how the blood microbiome changes when a bloodstream infection (BSI) occurs. Here, we analyzed the dynamics of the blood microbiome in a porcine model of polymicrobial bacteremia induced by fecal peritonitis. Serial blood samples were taken over 12 hours post-induction of fecal peritonitis, and BSI was validated by conventional blood culture and assessment of clinical symptoms.

**Results:** The bacterial populations in the blood microbiome were retained throughout the experimental period. However, there were significant taxonomic differences between the profile in the fecal and blood microbiomes, reflecting tropism for the blood environment. We also confirmed that the microbiota we detected was not contaminated by low mass bacteria in the bloodstream. However, at the same time, we noted a slight increase in *Bacteroidetes*, which is a major component of the gut microbiome, as sepsis developed. Comparison of the functional pathways in the blood and fecal microbiomes revealed upregulation of pathways involved in environmental interactions, and downregulation of those related to cell proliferation, in the former. Based on the enriched biological pathways, we concluded that communication and stress management pathways are essential for the survival of the blood microbiome under harsh conditions.

**Conclusion:** This study suggests that the microbiota can be stably retained in the bloodstream over time. Although further investigation in humans is required, we suggest that the blood microbiome may be another factor to be considered in the context of BSI and subsequent sepsis.

## Introduction

Bloodstream infection (BSI) is defined as a medical condition in which viable bacteria or fungi are present in the bloodstream [1]; BSI is a significant threat in human health, particularly as it can cause sepsis and organ dysfunction [2]. A survey of BSI incidence in America and Europe since the 1970s reported rates between 80 and 189 per 100,000 persons per year; this number has increased in recent years [3,4]. Furthermore, many cases progressed to critical conditions. BSI is estimated as the cause of 79,000–94,000 deaths per year in North America and 157,000 deaths per year in Europe [5].

Blood culture is a well-established method of detecting BSI, but the result is not always clinically relevant. A recent study reported that 42.6% of 2,659 of patients with suspected sepsis had positive blood culture results, whereas the remaining 1,526 patients (56.4%) were culture-negative [6]. False positives caused by contamination are a concern when testing for BSI. For example, only 51% of blood culture-positive samples are actual BSI (41% are due to contamination and 8% are of unknown clinical significance) [7,8]. The expected sensitivity and specificity of blood culture varies depending on the experiment conditions, which include collection time, skin preparation, sampling site, and sample volume [9].

High-throughput sequencing is an alternative technique for detecting microbes in the blood, even without culture [10]. However, this highly sensitive method raises other questions about BSI, i.e., is there a blood microbiome? (reviewed in [11]). Although the bloodstream is considered to be a sterile environment, recent evidence suggests that it may contain bacteria (or a microbiome), which may also colonize other organs. It was reported in 1969 that the metabolically active bacteria may be present in the blood [12], and recent studies propose that bacteria may use the bloodstream as a transport system. For example, bacteria have been identified in blood and adipose tissue samples from patients with type 2 diabetes [13] and in the liver of patients with non-alcoholic fatty liver disease [14]. Furthermore, *P. gingivitis* derived from chronic periodontitis infection is thought to contribute to Alzheimer’s disease [15], leading to speculation that the human microbiome can disseminate to other organs through the bloodstream.

However, most of these studies measured the blood microbiome at a single time point, making it hard to rule out the possibility of contamination. Suppose the microbiota is stably maintained in the bloodstream. In that case, it should be detectable over time, like other microbiome in the body, and concordance between time points can support the blood microbiome’s existence. However, this type of data is difficult to collect from human samples. Furthermore, these data should be assessed alongside blood culture results. Even in cases of BSI, a limited number of bacterial cells are present (about 0.1 to 100 bacterial cells per 1 mL of infected blood) [9]. If there is a microbiome in the bloodstream, it is questionable why those bacteria cannot be detected in the blood culture. So, a more controlled environment is required to evaluate the relationship between BSI and the blood microbiome.

Recently, we developed a porcine model of fecal-induced peritonitis [16]. We observed symptoms of organ dysfunction at about 7 hours (median) after introducing feces into the pig abdomen. The chance to detect bacteria in the blood culture also increased gradually after induction. Here, we used the same model to investigate the role of the blood microbiome in bacteremia. Surprisingly, many bacterial cells were detected in pig blood, even before fecal induction; it is likely that these bacteria comprise the blood microbiome. In addition, the bacterial species we identified in blood culture were not the dominant species detected by sequencing, although they presented similar tendency over times. The data presented herein provide new insight into the relationship between BSI and the blood microbiome.

## Materials and Methods

### Animal experiments

*Sus scrofa domesticus* (about 40 kg) were used for all experiments, as described previously [16]. Briefly, autologous feces were collected 1 day before the experiment and stabilized overnight at 4°C. Sepsis was induced by introducing feces (1 g of feces per 1 kg body weight, diluted in dextrose saline (10 g/dL) and heated to 37°C for 1 hour in a water bath) into the abdominal cavity of anesthetized animals. Blood samples (5 mL) were drawn from the intravenous line at 1- or 2-hour intervals and collected in DNA/RNA shield (Cat# R1150, Zymo Research, Irvine, CA, USA). All procedures were approved by SNUBH IACUC (BA1804-246/040-01).

### Capture of the blood microbiome

MBL-coated magnetic nanoparticles (MNPs; 2 mg/mL) were added to 3 mL of blood-DNA/RNA shield and incubated for 20 mins at room temperature. Captured bacteria were harvested using N52 magnets (BYO88-N52, KJ Magnetics, PA, USA) and washed with PBS to remove other blood components. Captured bacteria were stored in DNA/RNA shield at −20°C before use.

### Fecal sample preparation

Autologous feces (3 mL) from each pig were diluted with dextrose saline (10 g/mL) in a DNA/RNA Shield tube (Cat # R1150, Zymo Research, Irvine, CA, USA) and stored at −20°C before the experiment. Total genomic DNA was extracted using an MoBio PowerFecal® DNA Isolation Kit (Cat # 12830-50, MO BIO Laboratories, CA, USA) and FastPrep-24™ (MP Biomedicals, LLC, Irvine, CA, USA).

### Extraction of DNA extraction from the ZymoBIOMICS Microbial Community Standard

Total genomic DNA was extracted from 20 μL of ZymoBIOMICS Microbial Community Standard (Cat # D6300, Zymo Research) using a ZymoBIOMICS DNA Miniprep Kit (Cat # D4300, Zymo Research).

### Library preparation

Bacteria-captured beads were incubated at 37°C for 1 hour with 10 mg/mL of lysozyme (Cat # 10837059001, Roche, Basel, Switzerland) diluted in 10 mM Tris-HCl (pH 8.0) buffer. Next, the beads were transferred to lysis buffer (10 mM Tris-HCl (pH 7.4), 10 mM EDTA, and 2% SDS) containing 0.5 mg/mL proteinase K (Cat # B-2008, GeNetBio, Daejeon, Republic of Korea), and incubated overnight at 37°C. Genomic DNA was extracted from cell lysates by addition of PCI (phenol-chloroform-isoamyl alcohol) followed by overnight incubation at −20°C. Next, DNA was precipitated with 0.6 volumes of isopropanol and 0.1 volume of 3 M sodium acetate (Cat # SR2006-050-55, Biosesang, Seongnam, Republic of Korea). After washing the pellet with 70% ethanol and dissolving it for 30 min at 37°C in 100 μL of TE buffer, 1 μL of RNase A (Cat # B-2007, GeNetBio) was added for 30 min at 37°C to remove RNA contamination. Zymo DNA Clean & Concentrator-5 (Cat # D4014, Zymo Research) was used to purify DNA, which was eluted with 20 μL of RNase-free water.

### Sequencing of 16S rRNA V34 region

Because all samples (except the fecal samples) contained a limited amount of DNA, Q-RT-PCR was performed using MIC qPCR (Bio Molecular Systems, Upper Coomera, QLD, Australia) to determine the number of amplification cycles before saturation (typically 20–28 cycles). After running the initial PCR with V34 primers, eight cycles of adapter ligation PCR were performed. The final library was sequenced using Illumina MiSeq (Illumina, San Diego, CA, USA) with a 2 × 250 bp configuration. The raw sequencing data were deposited in the European Nucleotide Archive (ENA) under accession ID PRJEB39083.

### Taxonomy analysis

PCR amplicons amplified by the Illumina V34 PCR primers were selected using ipcress (provided in exonerate version 2.2) after concatenating paired-end reads [17]. Next, taxonomy information was assigned to each paired read using the RDP classifier (version 2.11) [18] and the RDP database (release 11.5) [19], with a confidence score > 0.8. The cut-off was determined by ZymoBIOMICS Microbial Community Standard sequencing as described in Figure S1. To define the operational taxonomic units (OTUs), concatenated V34 amplicons with 99% identity were clustered using VSEARCH (version 2.13.6) [20]. Ambiguous clusters were removed, and reads were included if they contained > 5% of reads assigned to a different genus when compared with the seed genus assigned to the same cluster. OTUs with low abundance (less than 0.4167% of OTUs) in all six samples were not entered into the taxonomic-based analysis. After taxonomic-level clusters were defined, additional levels of taxonomy (from genus to phylum [including family, order, and class]) were included. To analyze OTUs not observed at the initial time point (i.e., before induction of fecal peritonitis), clusters with no reads in the initial sample were selected. SourceTracker2 was used to analyze changes in the initial microbiome after feces introduction [21]. Default options were selected to run the program.

### Pathway analysis using PICRUSt2

To investigate differences in biological pathways among different microbiome populations, PICRUSt2 (version 2.2.0-b) [22] was used to estimate relative enrichment of biological pathways in each sample. KEGG orthologs and pathways inferred from the genus level (99% OTUs) were entered into the PICRUSt2 metagenome pipeline. KEGG ortholog results were transferred manually to KEGG pathways based on PICRUSt2-appended default files. KEGG ortholog enrichment values without pathway information were discarded and the remaining values were summed. If KEGG orthologs belonged to more than two KEGG pathways, they were added individually.

## Results

### The Blood Microbiome in the porcine model before induction of peritonitis

Because the number of bacteria in the blood is estimated to be 0.1–100 CFU/mL, even in BSI [9], we expected to observed similar numbers in the blood microbiome of the porcine bacteremia model. First, we enriched the number of bacterial cells in each 3 mL blood sample using opsonin-coated magnetic beads [23] and then extracted genomic DNAs prior to 16S rRNA gene sequencing using Illumina V4 primers. Across 12 time points, we obtained 116,062.39 paired reads on average per sample from each of the six animals (median 102,654 reads; minimum 38,160 reads; maximum 333,130 reads; all reads are available at the ENA under accession ID PRJEB39083). We clustered these reads to define the OTUs and then analyzed the taxonomy information (from phylum to the genus) using the RDP classifier [18] (See Methods for more details).

Surprisingly, we observed many bacterial species at the initial time point, even before induction of fecal peritonitis. Furthermore, those populations did not change much over 12 hours (Figure 1, phylum level; Figure S2, genus level). Compared with the fecal microbiome, which contained *Firmicutes* and *Bacteroidetes* as the most abundant phylum, *Proteobacteria* was the most abundant phylum in the blood. Four out of six animals tested (P1120, P1126, P1211, P1219; Figure 1) maintained a relatively consistent microbiome population, with few perturbations (e.g., at 11 hours post-induction in P1120 and 6 hours post-induction in P1219). Two other animals (P1016 and P1103) showed some fluctuations in microbial population composition during the early induction stage, but they showed similar patterns after 4 hours.

**Figure 1.**
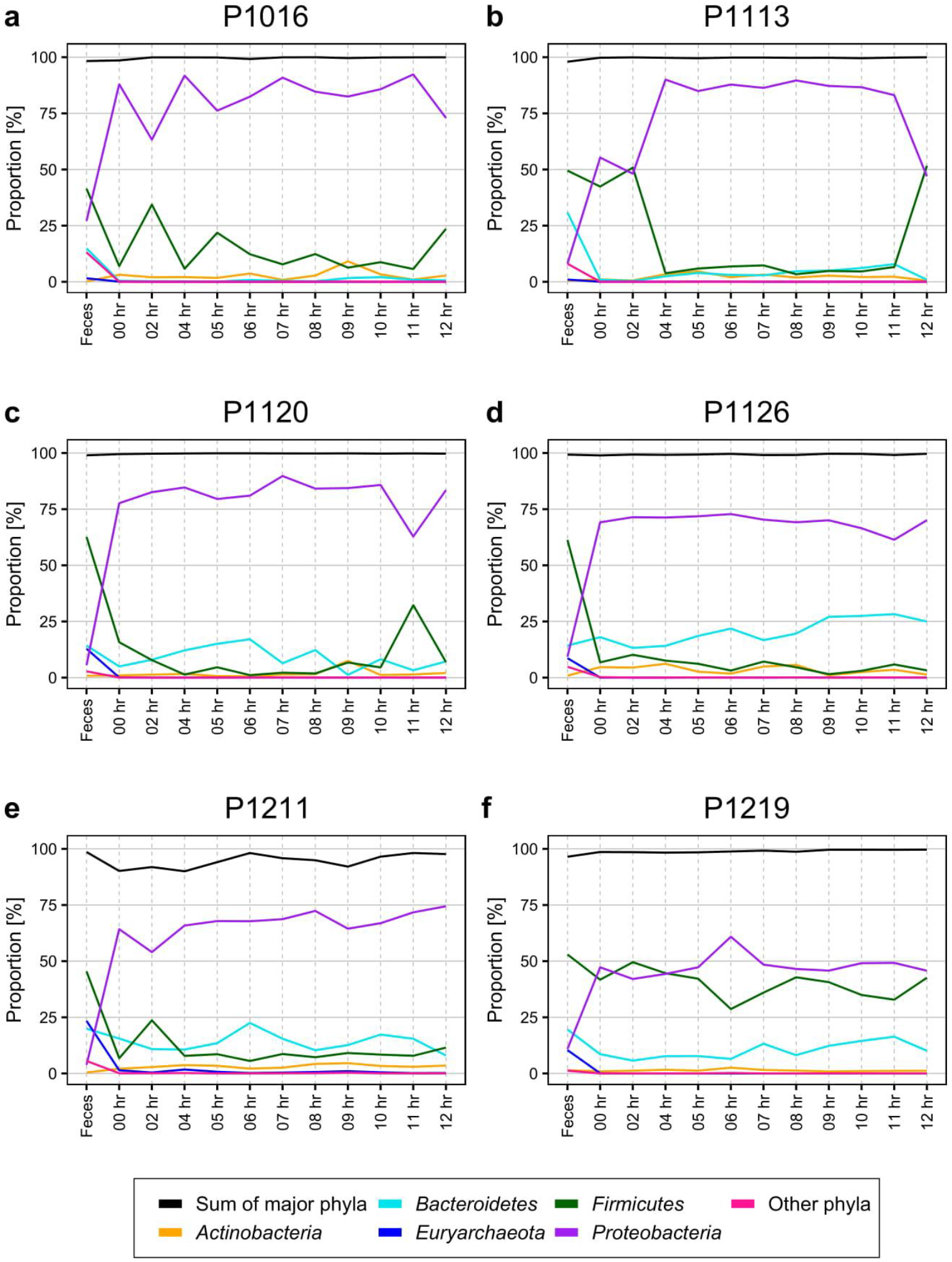
Percentage of Operational Taxonomic Units (OTUs) for each phylum identified in the blood of pigs with fecal-induced peritonitis. The proportion of each phylum in the blood microbiome was maintained consistently throughout the induction period, which is distinctive to that of the fecal microbiome. Although *Proteobacteria* was the major phylum identified in blood, the percentage varied in each of the six animals tested. The proportions at the genus level are shown in Supplementary Figure 2.

To confirm the presence of the microbiome before induction, we also performed a DNA-FISH experiment using the same bead-captured bacterial cells and a universal probe (5’-CTTGTACACACCGCCCGTCACACC-3’), which can hybridize to 98% of bacterial 16S rRNAs available in the GTDB (release 89) [24] (Fig. S3). Although the blood culture results for all samples were negative, we verified that blood samples obtained before peritonitis induction contained many bacteria. Taken together, the results suggest that the blood microbiome is maintained in the porcine bacteremia model, even before a positive blood culture result is obtained.

### Characterization of the initial blood microbiome

It is possible that the microbiome observed in the blood may be due to contamination of the arterial catheter used for sampling, or by the skin microbiome, as discussed previously [25–27]. Also, because the number of bacterial cells in our samples was small, the microbiome we observed could be caused by uncontrollable low biomass contamination from an unknown source, known as a ‘KitOme’ [28]. Suppose a majority of bacteria we identified were derived from any accidental contamination. In that case, we expect no discernible pattern on blood microbiome profiles. However, suppose bacteria from the unknown source were related to peritonitis induction. In that case, we may detect new types of bacteria to be introduced into the blood.

To test this hypothesis, we used SourceTracker2 [21] to compare the blood microbiome before and after peritonitis induction. We assumed that the initial time point would be the “sink” for contamination, and so attempted to identify trends in the ‘contamination’ over time. In all six animals, the bacterial population that detected at the initial time point (sink) decreased gradually up until about 6–8 hours after peritonitis induction, and then maintained a steady level up until 12 hours after induction (Figure 2). This trend matched the physiological symptoms of sepsis [16]; therefore, we speculated that the blood microbiome was altered about 6–8 hours after fecal induction. When we used 12-hour samples as sinks, we did not observe any trends (Fig. S4).

**Figure 2.**
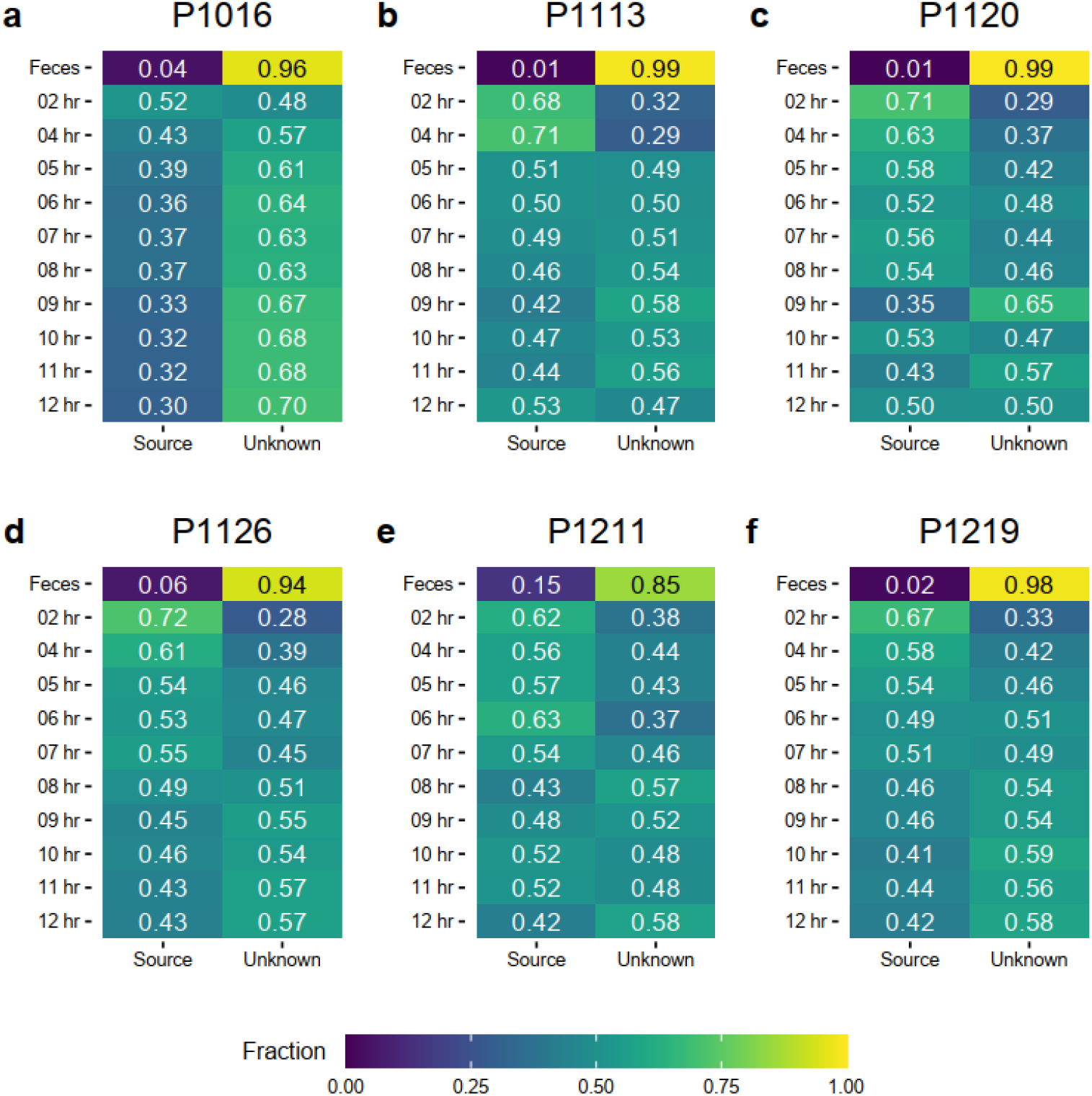
SourceTracker shows that the population of novel bacterial strains not present at the initial time point increased throughout the induction period. The microbiome observed at the initial time point was designated as the “source.” The proportions of species that originated from the source were then tracked throughout the induction period. The proportion of unknown strains increased gradually until about 7 to 8 hours after induction before being maintained.

To validate the result ruling out low biomass contamination, we also analyzed two public datasets: one was a serially-diluted mock community standard [29] and the other was a serial dilution of *S. bongori* [30] (Fig. S5). As expected, when we used the mock community standard sample as a sink, we observed a linear reduction in its proportion as the dilution factor increased. However, when we used the most dilute sample as a sink (mimicking low biomass at the initial time point), we observed no changes in the population. Similarly, in the *S. bongori* study [30], we used 10^8^ cells or 10^3^ cells as the sink, and found that the populations were different from those in the blood microbiome. We concluded that the blood microbiome we observed is due nether to contamination caused by sample preparation nor to “noise” artifacts (KitOme).

### A new blood microbiome was detected after peritonitis induction

Although SourceTracker2 analysis revealed noticeable changes in the blood microbiome after peritonitis induction, the composition of overall bacterial population was retained. We speculated that the pre-existing blood microbiome might mask small changes in the blood microbiome caused by peritonitis induction. To test this, we computationally discarded OTU clusters containing any OTUs observed at the initial time point, obtaining OTUs newly emerged at each sample. We found gradual changes in bacterial populations as BSI progressed (Fig. 3).

**Figure 3.**
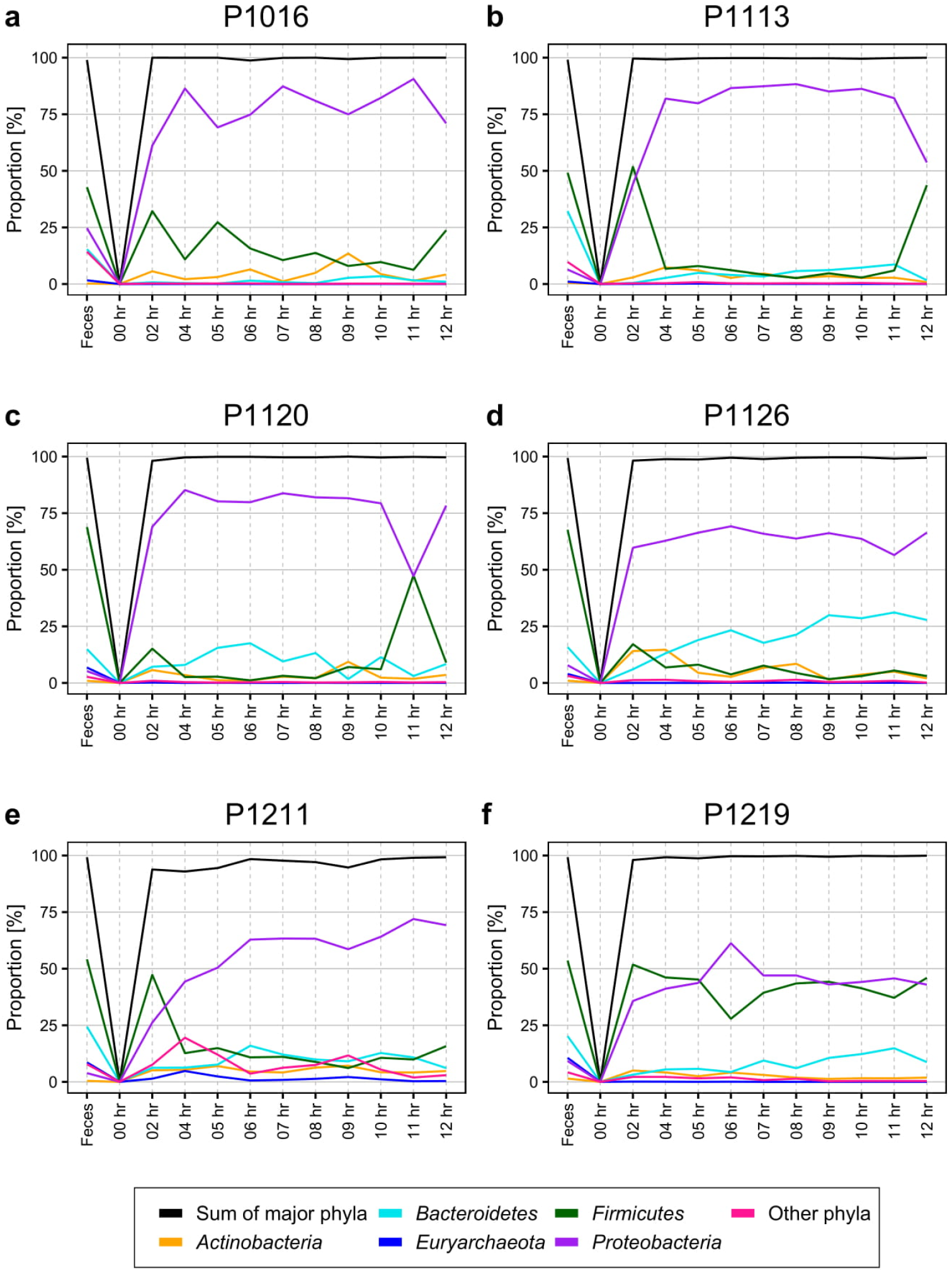
Percentage of newly emerged Operational Taxonomic Unit (OTU) for each phylum in the blood of pigs with fecal-induced peritonitis. The microbiome observed at the initial time point was discarded computationally from all other time points and the proportion at each time point was re-calculated.

At the phylum level, the blood and fecal microbiomes harbored different populations of *Proteobacteria* and *Firmicutes*. *Proteobacteria* is not dominant in feces, but it was the most abundant phylum in the bloodstream of all six animals. *Firmicutes* showed the opposite trend. This alteration happened regardless of computational filtering of the intrinsic microbiome population. Interestingly, the representative gut microbiome phylum *Bacteroidetes* [31] was detected at all time points and showed a gradual increase over time; this was not observed in the absence of the background blood microbiome (light blue line in Fig. 3). However, the composition of the blood microbiome did not change, even though we discarded the background populations (i.e., the microbiome detected before fecal induction). Even after introduction of new bacterial species post-induction, the composition of the blood microbiome did not change much.

### Comparison of results with those from blood culture and physiological data

As the standard diagnostic method for the BSI is blood culture, we also conducted clinical laboratory blood culture tests to identify bacteria causing septic symptoms after fecal induction. As reported previously [16], of 82 bacterial species identified by blood culture, 83% of the cases were *E. coli* (57.3%) or *Streptococcus* (25.7%) species. We also identified those species by sequencing and at similar percentages (Fig. 4). For example, *E. coli* and *S. dysgalactiae* were identified in animals P1016 and P1120 at the later stages after induction, and we also observed a significant increase in these species at the time with positive blood culture results. In animal P1126, *S. alactolyticus* was detected only at 7–8 hours after induction, and *E. coli* was detected at a later time point. Similarly, we observed a slight increase in *Streptococcus* at 7-8 hours after induction. *E. coli* was detected in animal P1219 at an early stage, and the sequencing data also supported a higher *E. coli* population throughout the experiment.

**Figure 4.**
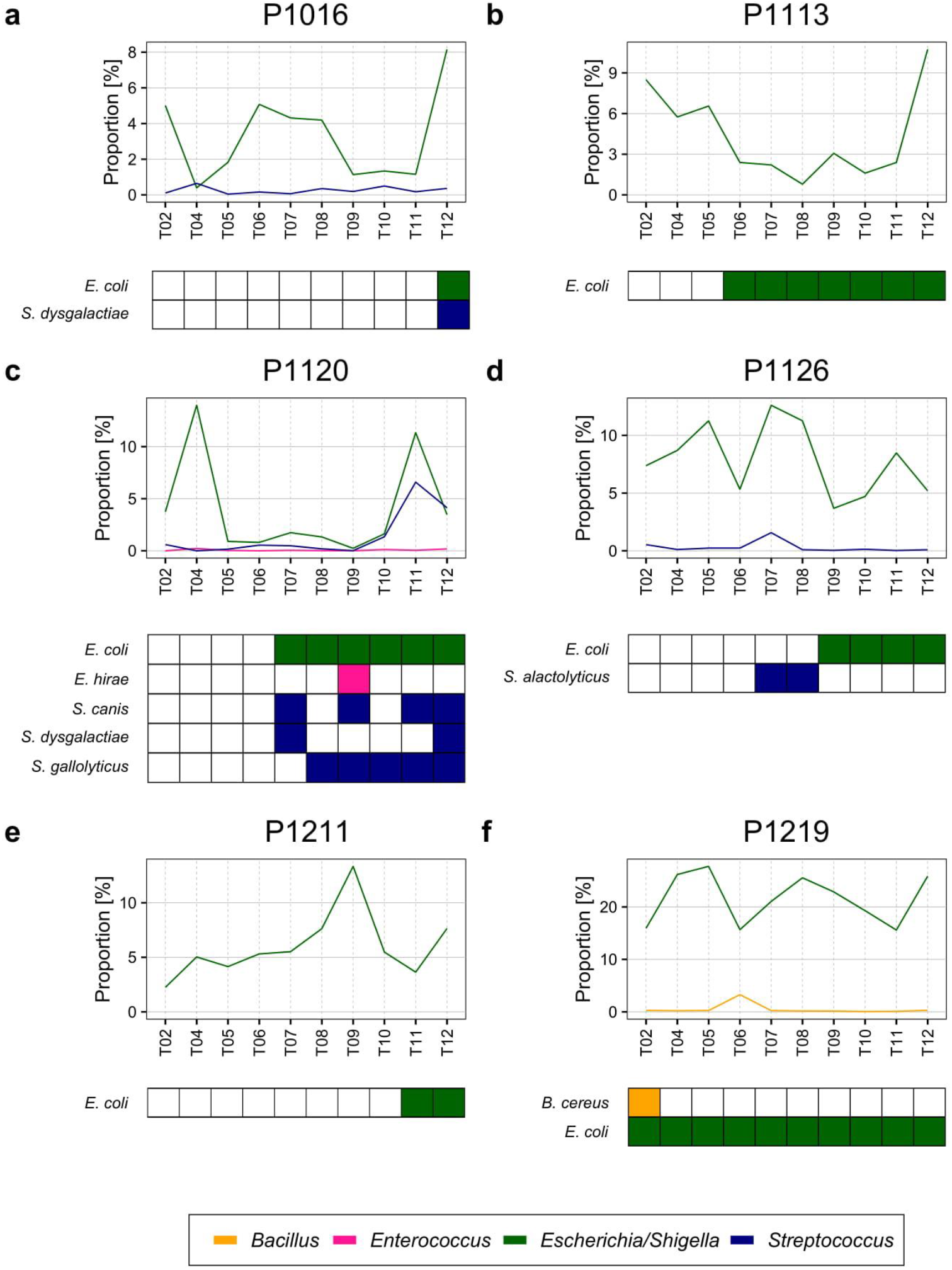
Bacteria detected in blood cultures. *E. coli* and other species (*S. dysgalactiae, E. hirae, S. alactolyticus, S. gallolyticus, B. cereus*) confirmed in blood cultures were also observed in our blood microbiome sequencing result from each sample, although their associations were moderate.

However, according to our microbiome profile data, bacterial species confirmed by blood culture were not the most abundant in the bloodstream, even 12 hours after peritonitis induction. One possible explanation for this is that bacterial DNA from white blood cells [32] or cell-free DNA [33,34] was detected by sequencing; these bacteria may be present at numbers too low to be detected by culture-based methods. Although microbial DNA could induce inflammation and other host responses, they are a false positive with respect to the blood microbiome. Another reason may be dormant bacteria, which cannot be cultured under test conditions [35–37]. Because dormant bacteria can reactivate depending on the environmental conditions, they could contribute to blood microbiome function even if they are not culturable immediately. Further investigation is required to explain these discrepancies.

Previously, we reported that all animals showed symptoms relevant to sepsis, which developed gradually 5–6 hours after fecal induction [16]. In addition to the blood culture results, we compared the emergence times of the altered blood microbiome with the timing of physiological symptoms (Fig. 5). Based on SourceTracker analysis, we found that the levels of cytokines (IL-1β and IL-6) in the blood increased as the new microbiome emerged (Fig. 2). Although we cannot tell whether this new blood microbiome plays a role in septic symptoms during the experiment, the data indicate that the blood microbiome might have some function(s).

**Figure 5.**
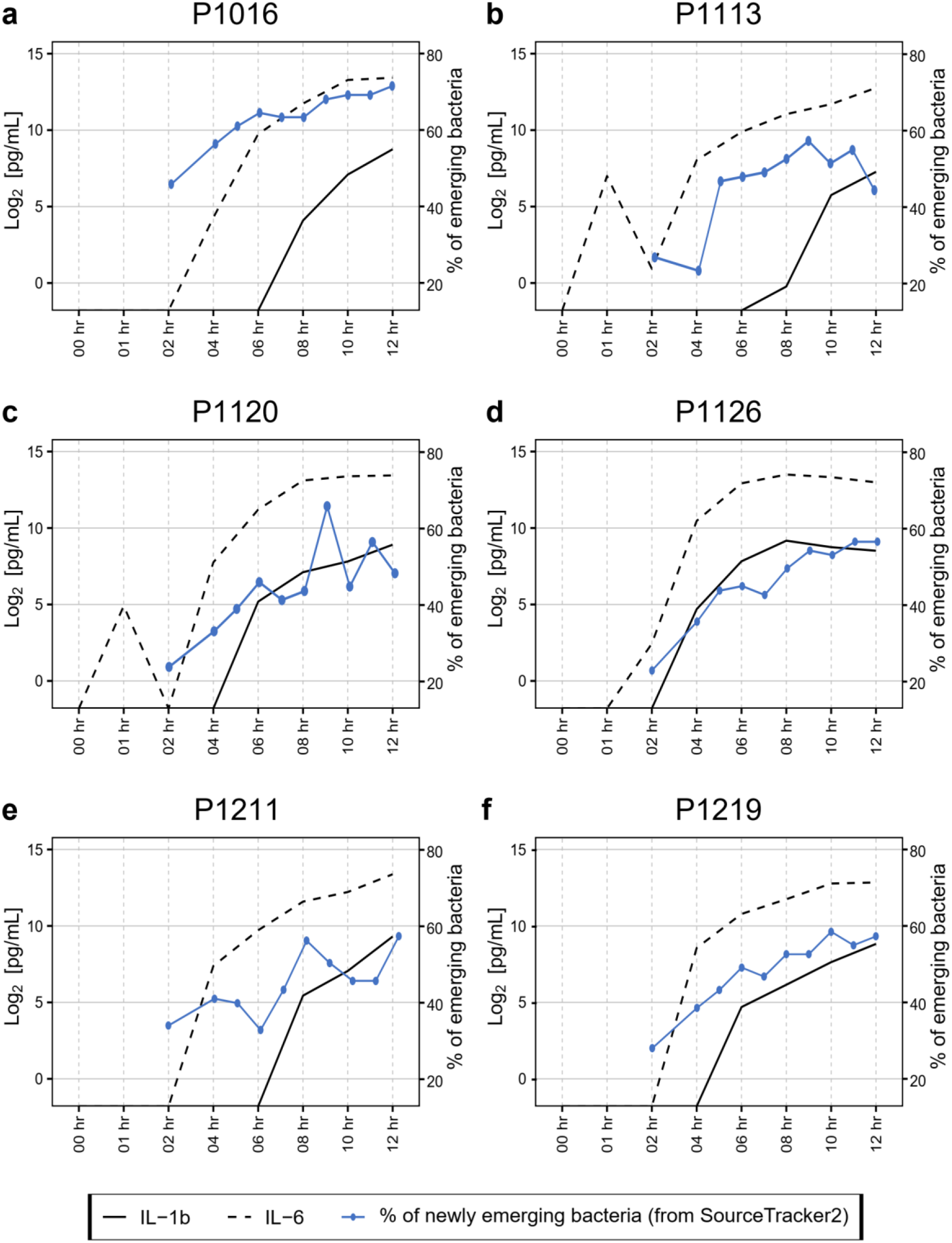
Association between cytokine induction and introduction of new bacterial species into the bloodstream. Measurement of IL1β and IL6 levels in the blood showed that most animals began to develop septic responses between 4 and 8 hours after induction. The proportion of newly emerging bacteria was measured by SourceTracker2 (Fig. 2) by comparing the altered microbiome with the microbiome detected before peritonitis induction. The original cytokine data were published previously [16].

### Biological pathways favored in the blood microbiome

Genes and pathways enriched in a particular population are more relevant to function than bacterial composition [38,39]. The blood of the porcine bacteremia model used here is a unique environment for bacteria, which contains high levels of inflammatory cytokines, immune cells, and a unique composition with respect to chemicals such as lactic acid [16]. Therefore, we performed PICRUSt2 analysis to identify putative functions of the blood microbiome (Fig. 6). We found that pathways related to the ABC transporter, the two-component system, and oxidative phosphorylation were enriched in the blood microbiome. By contrast, pathways related to purine metabolism, pyrimidine metabolism, and ribosome expression were downregulated compared with those in feces.

**Figure 6.**
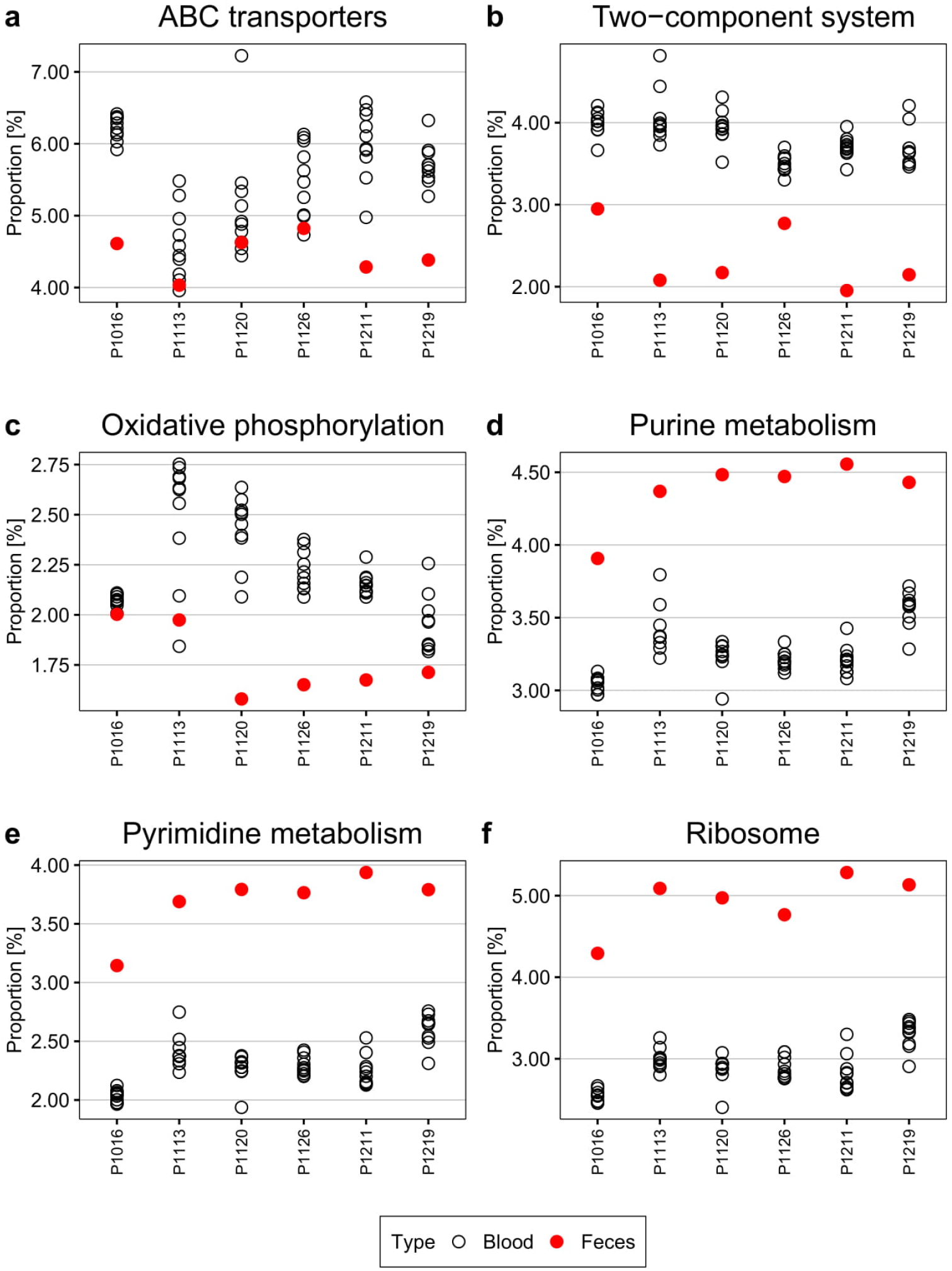
KEGG pathways differentially presented by the blood microbiome. Pathways related to ABC transporters, the two-component system, and oxidative phosphorylation were over-represented in the blood microbiome compared with the fecal microbiome. By contrast, pathways related to central metabolism, such as purine and pyrimidine metabolism, and ribosome-related pathways, were under-represented.

The ABC transporter and the two-component system are essential components that allow bacteria to respond appropriately to environmental signals [40–43]. For example, some bacteria regulate acid-base balance and metal iron homeostasis by utilizing ABC transporters [40,41]. Furthermore, bacteria can use this transporter to defend themselves against antimicrobial peptides and proteins in the bloodstream [40,42]. The two-component system, which is composed of a histidine kinase and a response regulator, is a major bacterial central signaling pathways that senses environmental cues [43,44] such as pH [45,46] and oxidative stress [47]. It is also tightly linked to bacterial responses to the host immune system [48,49]; therefore, it is likely that cells in the microbiome would activate to survive.

Conversely, purine metabolism, pyrimidine metabolism, and ribosome expression were suppressed in the blood microbiome, which may limit cell proliferation and growth [50–52]. Because the bloodstream is a harsh environment, cells may downregulate essential metabolic functions to survive. Simultaneously, reducing cell proliferation or maintaining low metabolic activity (e.g., dormancy) may enable bacteria to escape immune surveillance in the blood [35,36].

## Discussion

Here we analyzed the blood microbiome of pigs with bacteremia to investigate its role in BSI. Previous studies report microbiota transmission among organs, presumably through the bloodstream [13–15]. However, it is unclear how the blood microbiome is maintained because these studies provide only a ‘snapshot’ at a single point in time. By monitoring the blood microbiome over time, we confirmed that each animal retained a relatively consistent blood microbial population. Furthermore, by identifying potential pathways enriched in this population, we revealed that bloodstream bacteria might be adapted to respond to environmental through use of ABC transporters and the two-component system. On the other hand, bacteria may not to grow under these unfavorable conditions, so pathways related to nucleotide biosynthesis may be suppressed.

The biggest challenge to monitoring bacteria in the bloodstream was low numbers in comparison to the microbiome at other body sites; such low numbers mean that even minor contamination can have a significant effect on detection results [29,53]. To overcome this, we selectively enriched bacteria from blood using opsonin-coated MNPs [23] and identified them by conventional 16S rRNA gene sequencing. Still, it may be difficult to distinguish actual bacteria from bacterial DNA derived from the debris circulating in the bloodstream because the sequencing-based method is destructive. Therefore, after capturing bacteria with MNPs, we also performed RNA FISH to target the common region within the bacterial 16S rRNA gene; this confirmed that bacteria were present in the bloodstream, even before induction of peritonitis (Supplementary Fig. S3).

The most common source of contamination of the blood microbiota study is the skin microbiome. One study reported that *Firmicutes* (55.6%), *Bacteroidetes* (20.8%), *Actinobacteria* (13.3%), and *Proteobacteria* (5.1%) are representative phyla resident in the porcine skin microbiota [54]. Among them, *Staphylococcus* is a dominant genus within the skin microbiota of animals [55] and humans [56,57]. However, these bacteria were not a major component of the blood microbiome population. Because we drew blood via a catheter, we should have observed skin contamination, even in blood cultures. Furthermore, when we filtered identical bacterial sequences detected at the initial time point, the newly emerged microbiota still had a similar composition in terms of bacteria populations (Fig. 3). Thus, we concluded that the blood microbiome was not contaminated by the skin microbiota.

The bacteremia model used herein had detectable bacteria in the blood after induction (confirmed by the culture test) [16]. Thus, we used this as a “positive control” to detect the blood microbiome. We observed that the composition of the blood microbiome changed gradually after induction of peritonitis by autologous feces, and that over-representation of *Bacteroidetes* in the gut microbiota [31] increased slightly over time (Fig. 3). The gut microbiome can enter the bloodstream when the host is immunocompromised [58]; thus, this may be a septic symptom of peritonitis. When we systematically traced the source of the microbiota, we also found that new populations were introduced gradually to the bloodstream after 4–6 hours of peritonitis induction (Fig. 2), mirroring clinical symptoms such as increased cytokine production (Fig. 4). However, it is not clear whether these newly introduced bacteria induce the septic symptoms or entered the blood because of sepsis.

Surprisingly, the bacterial species identified in the standard clinical laboratory culture tests were not the dominant species identified by our culture-free bacterial population analysis, although there was a moderate association (Fig. 5). We speculate that culturable bacteria comprise only a small portion of the total blood microbiota. Also, the blood microbiome may harbor a type of homeostatic mechanism that maintains the population. Like the commensal microbiome that maintains its population when exposed to exogenous bacteria [59–61], the blood microbiome may have a similar protective mechanism. However, because the blood microbiome may not be metabolically active relative to other commensal bacteria, further study would be required to examine this possibility.

The porcine peritonitis model we used here provides a unique opportunity to study the blood microbiome. Due to the large body size of pigs, we can monitor the dynamics of the blood microbiome over time by multiple sampling, which is challenging when using a small animal model such as a mouse. Although the animals we used for the experiment were not raised in an aseptic environment, the multiple sampling approaches and computational methods for population comparison used herein made it possible to characterize the blood microbiota comprehensively. Thus, this method may provide an excellent platform for developing new diagnostic techniques to detect BSI [10,27,33].

## Conclusions

Here, we reported changes in the composition of the blood microbiome in a porcine bacteremia model. By analyzing the blood microbiome in the same individual animal over time, we showed that it remains relatively consistent, even after induction of peritonitis. However, at the same time we found that new bacterial populations enter the bloodstream, the dynamic patterns of which are similar to those observed during a physiological response to BSI (e.g., cytokine elevation). By analyzing population-based pathways, we also confirmed that sensing mechanisms such as the ABC transporter and the two-component system are upregulated in the blood microbiome. Conversely, central nucleotide metabolism, which is essential for cell proliferation and growth, is suppressed, which probably helps the blood microbiome to survive the harsh bloodstream environment and escape immune surveillance. Our study also suggests that further investigations of the blood microbiome are required to improve current diagnostic approaches to BSI.

## List of abbreviations

ABC transporter: ATP-binding cassette transporters
BSI: Bloodstream infection
FISH: Fluorescent in-situ hybridization
KEGG: Kyoto Encyclopedia of Genes and Genomes
NMP: Magnetic nanoparticle
OTU: Operational Taxonomic Unit
PCR: Polymeric chain reaction
PICRUSt: Phylogenetic Investigation of Communities by Reconstruction of Unobserved States

## Declarations

### Ethical approval

All the procedures used for animal experiments were approved by SNUBH IACUC (BA1804-246/040-01).

### Consent for publication

Not applicable.

### Availability of data and materials

The raw sequencing data generated during the current study have been deposited in the European Nucleotide Archive (ENA) under accession ID PRJEB39083.

### Authors’ contributions

HH performed the microbiome experiments and data analysis, assisted by SMY and HSK; MSL prepared the blood microbiome samples and performed the FISH experiment; IP performed the animal experiments, assisted by DHJ and SK; HK, JHK, JHL, and TK conceived the study and designed the experiments; JHK, JHL, and TK analyzed the data; all authors helped to write the manuscript. All authors read and approved the final version.

## Acknowledgments

We thank You Hwan Jo, Doyun Kim, Hyunglan Chang, and Hyuksool Kwon from the Seoul National University Bundang Hospital for helping with the animal experiments.

---

**Figure S1.**
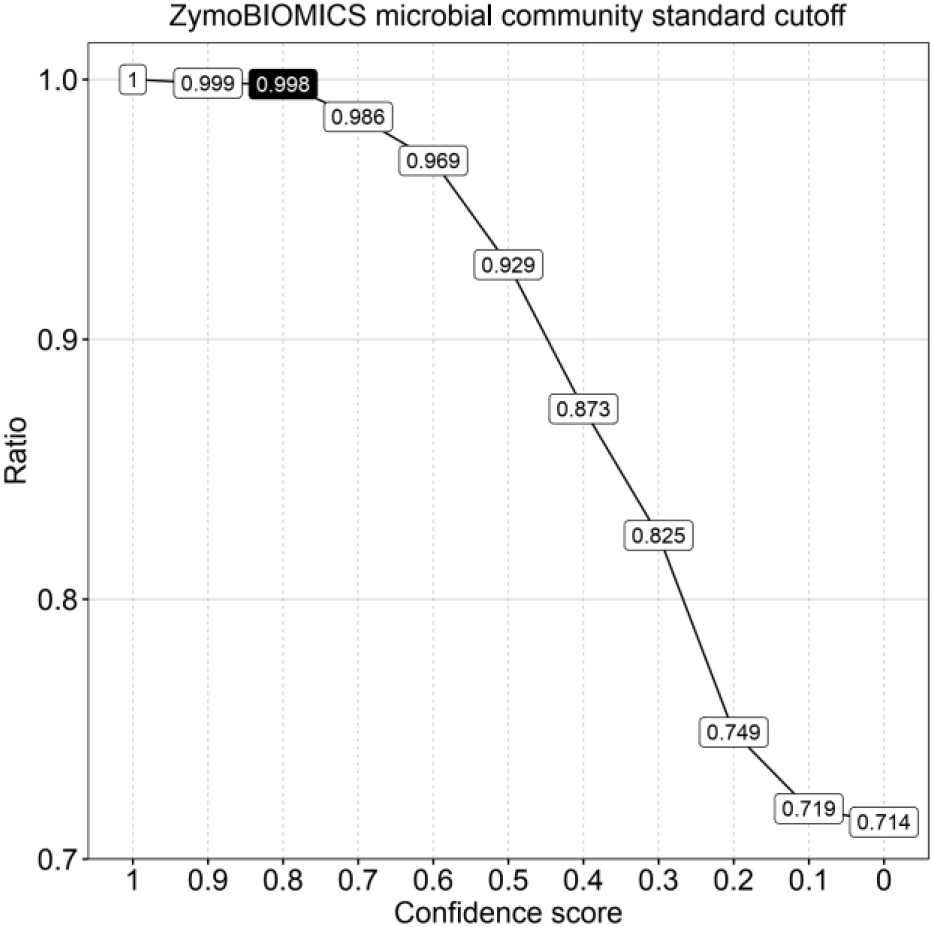
The cut-off value of the RDP classifier, based on standard microbiome data. The V34 amplicon of 16S rRNA was sequenced using the Zymo Microbiome Standard sample, which contains eight different bacterial genomes. The data were mapped to the GreenGene database using the RDP classifier (version 2.11) and the number of true positive reads (numbers inside the box) was evaluated. Based on this, the RDP classifier confidence score was set as 0.8 (cut-off for analysis), which assigned 99.8% reads correctly to their target.

**Figure S2.**
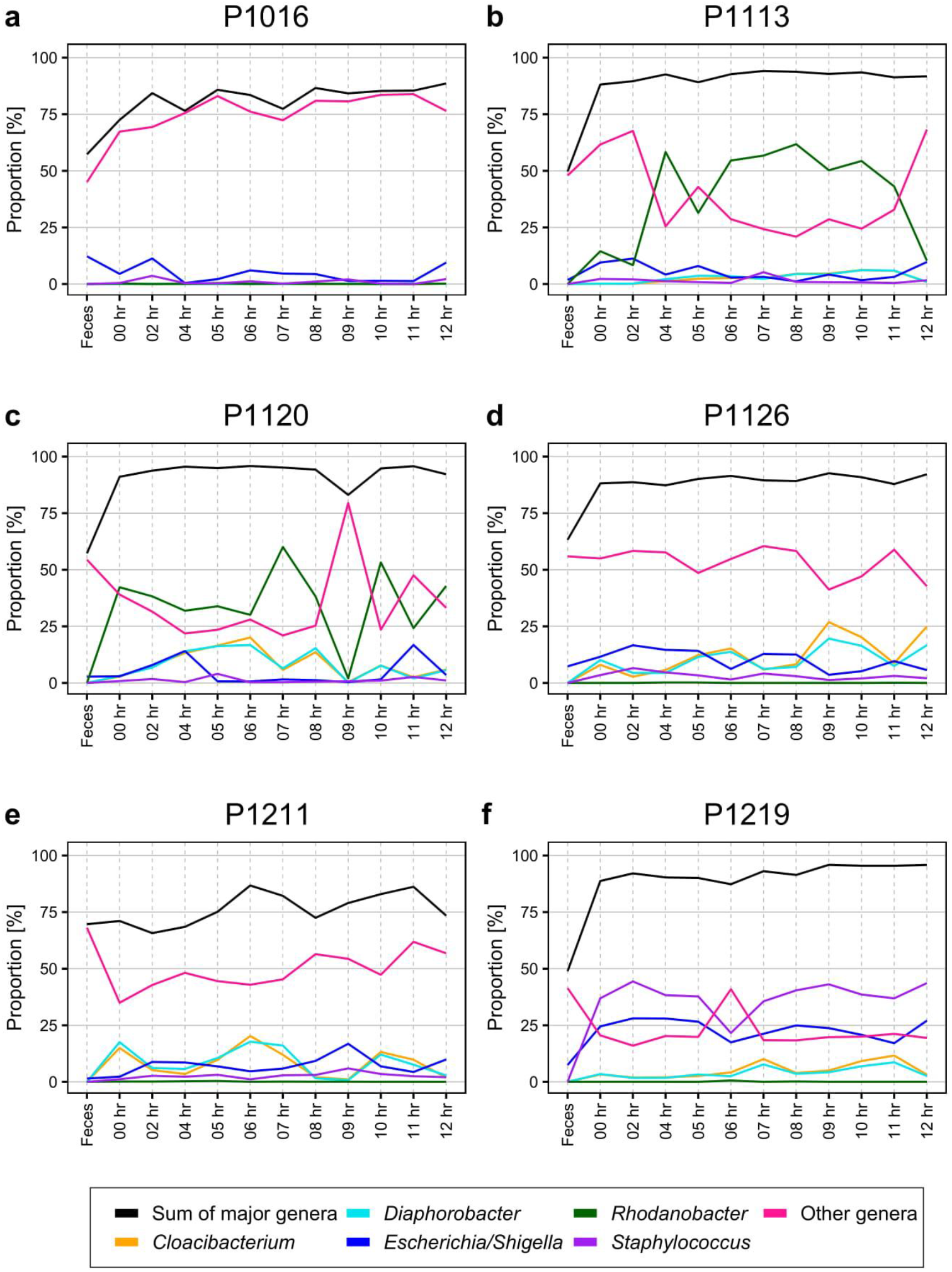
Percentage of Operational Taxonomic Unit (OTU) for each genus identified in the blood of pigs with fecal-induced peritonitis.

**Figure S3.**
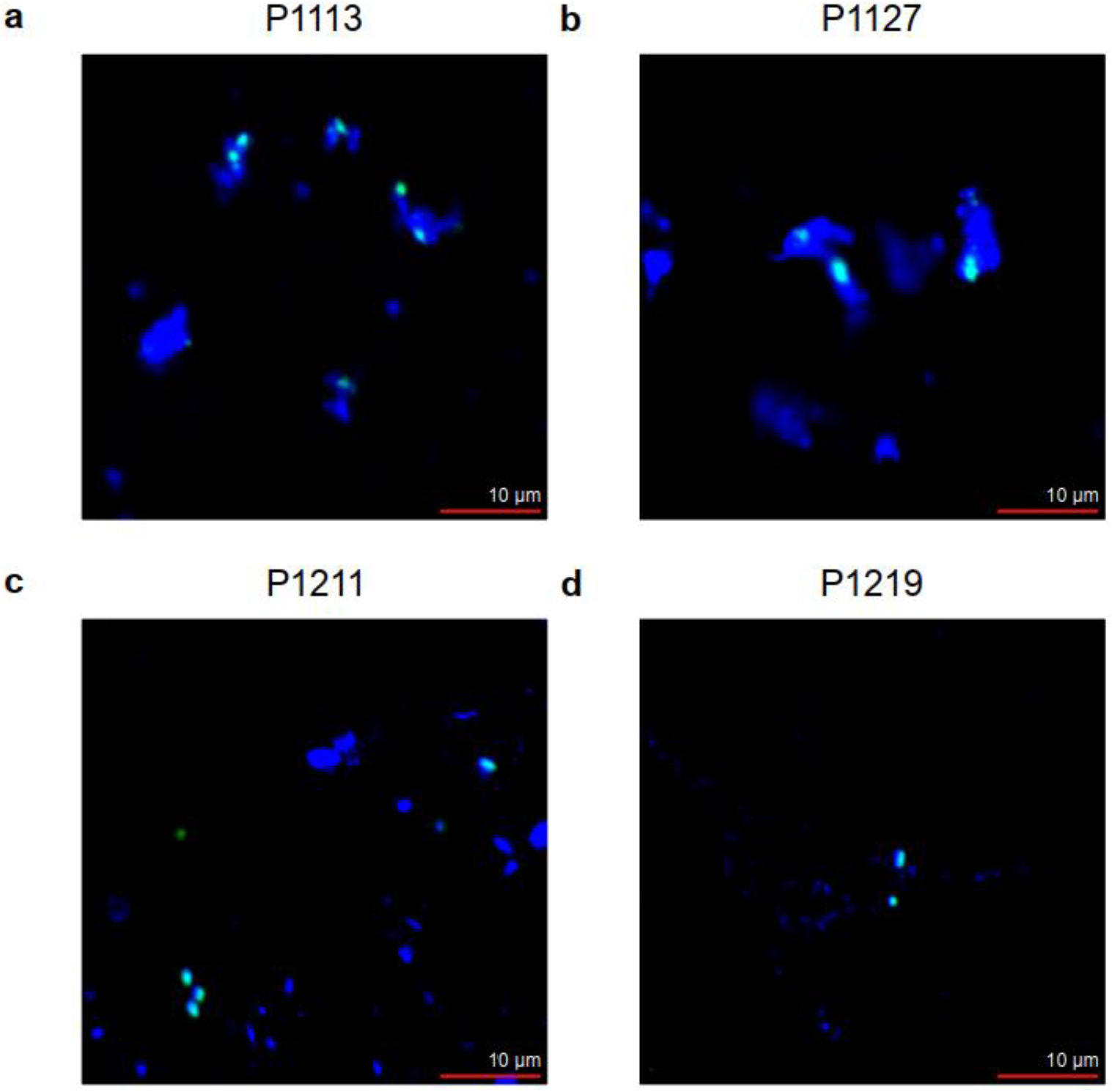
RNA-FISH results before fecal induction. Because blood cells are also captured, larger DAPI signals (blue) were also observed. We used ‘the common probe’, which targets the conserved region of 16S rRNA, to detect bacteria in the bloodstream. The green signal overlapped with the blue (cyan in this merged image), thereby highlighting the blood microbiome.

**Figure S4.**
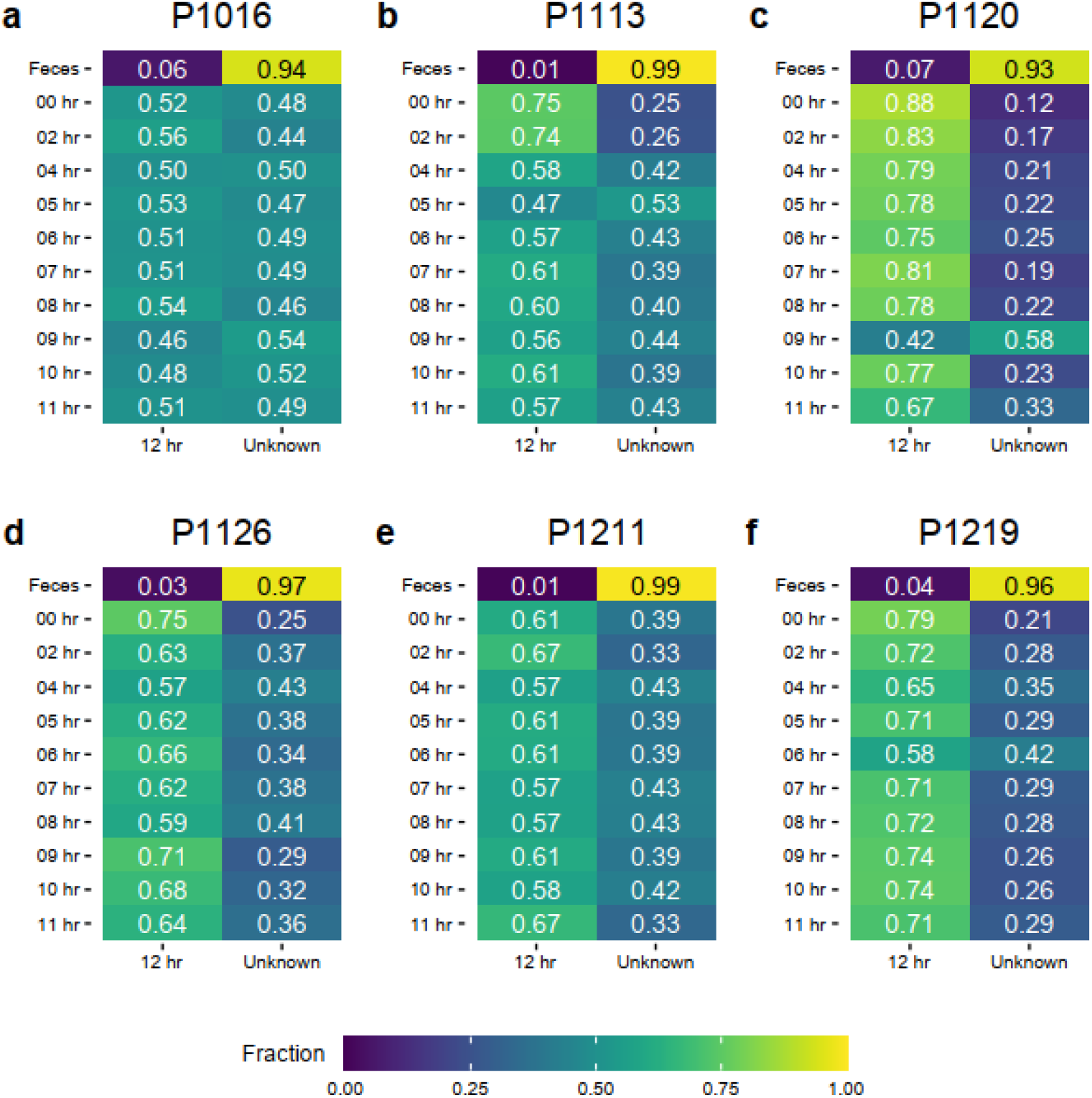
SourceTracker shows random patterns when the microbiome observed at the last time point (12 hours after peritonitis induction) was used as the source. Almost all samples contained more than 50% of the bacterial species identified at the end of the experiment, but those numbers did not change during peritonitis induction. Thus, no time-related changes were observed (compared with Fig. 2).

**Figure S5.**
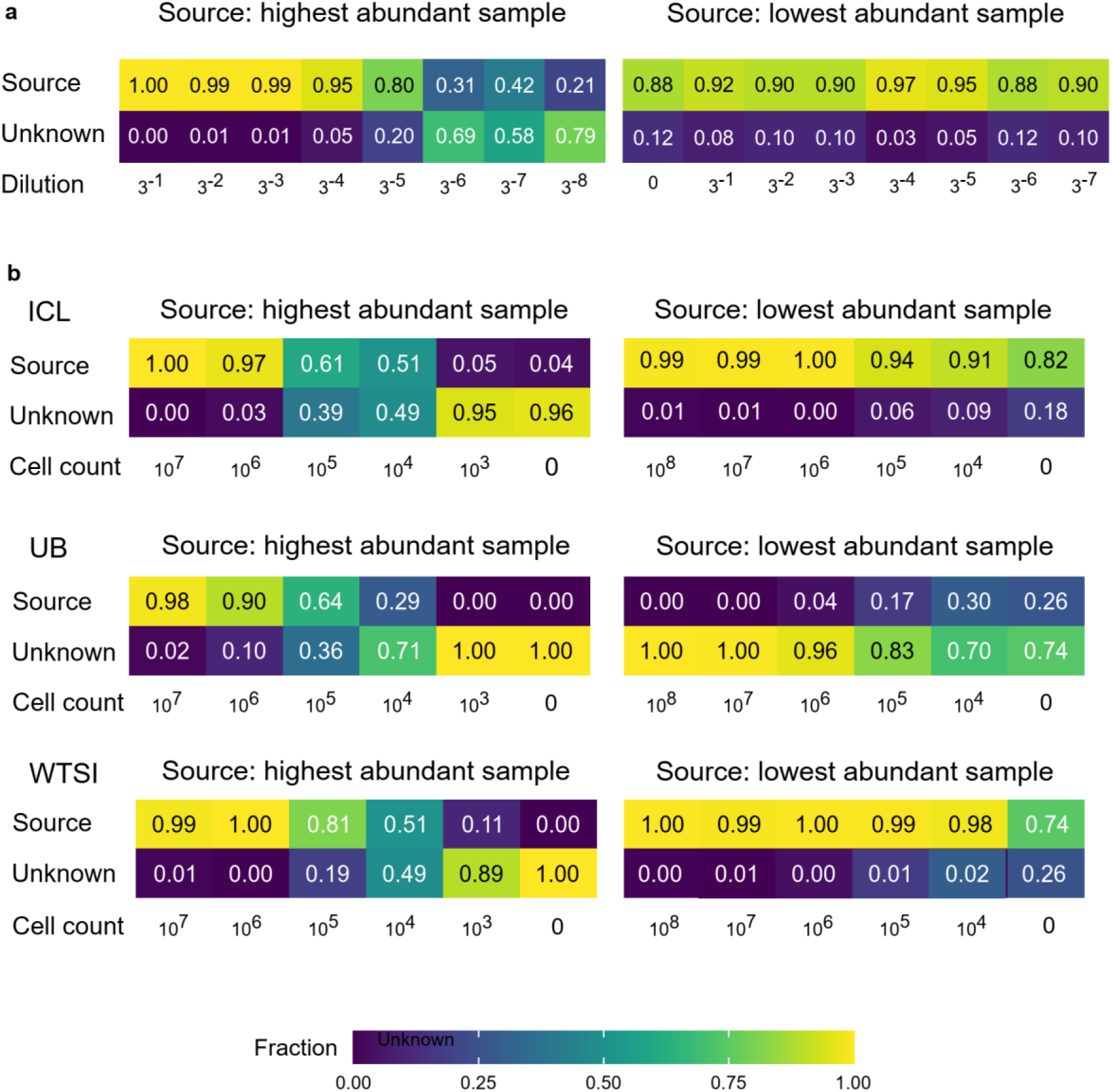
SourceTracker shows distinctive patterns for the contamination due to low abundant cell numbers in the sample. Serial dilution of the sample with the highest abundance showed a gradual decrease in the proportion of the original population (left) in all samples. However, when we used the low abundance sample (3^−8^ dilution for (a) or 10^3^ cells for (b)) as the source, no pattern was observed.

